# Hormones do not make the mole-rat: no steroid hormone signatures of subordinate behavioral phenotypes

**DOI:** 10.1101/2022.04.14.487730

**Authors:** Ilapreet Toor, Mariela Faykoo-Martinez, Phoebe D. Edwards, Rudy Boonstra, Melissa M. Holmes

**Affiliations:** Ecology and Evolutionary Biology, University of Toronto, Toronto, ON; Cell and Systems Biology, University of Toronto, Toronto, ON; Psychology, University of Toronto Mississauga, Mississauga, ON

**Keywords:** androgens, behavioral phenotype, cooperative breeding, DHEA, estradiol, naked mole-rat, progesterone, social behavior, social status, testosterone

## Abstract

In some cooperatively breeding groups, individuals have distinct behavioral characteristics that are often stable and predictable across time. However, in others, like the eusocial naked mole-rat, evidence for behavioral phenotypes is ambiguous. Here, we study whether the naked mole-rat can be divided into discrete phenotypes and if circulating hormone levels underpin these differences. Naked mole-rat colonies consist of a single breeding female and dozens to hundreds of non-reproductive subordinates. The subordinates can potentially be divided into soldiers, who defend the colony; workers, who maintain it; and dispersers, who want to leave it. We established six colonies de novo, tracked them over three years, and assessed the behavior and hormone levels of the subordinates. We found that soldiers tended to be from earlier litters and were higher ranked compared to workers, whereas dispersers were distributed throughout litters and rankings. There was no difference in estradiol, testosterone, or dehydroepiandrosterone (DHEA) levels amongst phenotypes. Progesterone levels were higher in soldiers but this difference appeared to be driven by a few individuals. Principal component analysis demonstrated that soldiers separated into a discrete category relative to workers/dispersers, with the highest ranked loadings being age, weight, and testosterone levels. However, the higher testosterone in soldiers was correlated with large body size instead of strictly behavioral phenotype. Workers and dispersers have more overlap with each other and no hormonal differences. Thus the behavioral variation in subordinate naked mole-rats is likely not driven by circulating steroid hormone levels but rather stems from alternative neural and/or neuroendocrine mechanisms.

## Introduction

Cooperative group living has evolved independently in numerous taxa. Within some of these groups, individuals have distinct behavioral characteristics that are often stable and predictable across time, or “behavioral phenotypes.” For example, cooperatively breeding male banded mongooses show consistent individual differences throughout their lifetime in their contributions towards cooperative and reproductive behavior (Sanderson et al., 2015). Likewise, female meerkats also show - consistencies in their levels of helping behaviors, such as babysitting and provisioning, over their lifetime (Carter et al., 2014). Cooperative group dwellers can also exhibit age-related polyethisms, where aging shifts an individual’s social role in the colony. For example, the eusocial Damaraland mole-rat (*Fukomys damarensis*) exhibits size and age-based polyethisms in cooperative behavior, and individuals have plasticity in their levels of cooperative behavior over time (Zöttl et al., 2016). In another eusocial mammal, the naked mole-rat (*Heterocephalus glaber*), in which eusociality is thought to have evolved independently, evidence for the existence of polyethisms and behavioral phenotypes remains equivocal.

Naked mole-rats are rodents native to East Africa that reside in large subterranean colonies with the largest reproductive skew reported in mammals (Brett, 1991; Jarvis, 1981). Colonies consist of one reproductive queen, 1-3 breeding male consorts, and dozens to hundreds of socially suppressed and non-reproductive subordinates (Faulkes et al., 1990b; Jarvis, 1981). It is estimated that most individuals spend their entire lives without reproducing (Jarvis et al., 1994). Thus, their fitness benefits are largely through their parents, and colony success and cooperation are key to survival. Naked mole-rat subordinates exhibit individual differences in their levels of cooperative behavior, but it is unclear whether differences are discrete and fixed phenotypes or a behavioral spectrum (Gilbert et al., 2020; Holmes and Goldman, 2021). Historically, subordinates have been described as either smaller “frequent” and “infrequent workers” or larger “non-workers” (Jarvis, 1981). Subsequent studies classify the larger individuals as soldiers since they are particularly active in colony defense against unfamiliar conspecifics and predators (Lacey and Sherman, 1991). This defensive behavior is distributed bimodally among subordinates, whereas working behavior is distributed on a spectrum (Mooney et al., 2015). A third phenotype that is dispersive is also described as individuals who have a strong drive to leave the colony, in contrast to their highly neophobic and xenophobic colony mates (O’Riain and Jarvis, 1997; O’Riain et al., 1996). The dispersers prefer contact with unfamiliar conspecifics and are generally larger colony members of both sexes (O’Riain et al., 1996; Braude, 2000; Toor et al., 2020; Toor et al., submitted). Historically, they are described as having higher levels of luteinizing hormone than other subordinates (i.e., closer to reproduction), show little working behavior, and have extensive fat reserves (O’Riain et al., 1996). Female dispersers also exhibit higher levels of within-colony aggression than other subordinates though, by definition, show lower aggression towards unfamiliar animals (Toor et al., 2020).

Some evidence exists for task specialization and task switching within the colony when specialist individuals are removed (Mooney et al., 2015), but recent reports also find a lack of specialization for individual cooperative behaviors (Siegmann et al., 2021). However, individuals who engage in one cooperative behavior are likely to engage in others as well (Siegmann et al., 2021). Furthermore, it seems there are some age-related polyethisms in subordinate levels of working behavior, but this plateaus after about two years of age (Gilbert et al., 2020). Recent reports reject task specialization in the cooperatively breeding naked and Damaraland mole-rats; however, these studies quantify specialization amongst types of working behavior only (digging, food carrying, nest building) and do not assess defense and dispersive behaviors in their analysis (Siegmann et al., 2021; Gilbert et al., 2020; Thorley et al., 2018).

Regardless of whether these behavioral profiles are truly categorical and stable, it is important to understand the physiological underpinnings of behavioral variation between individuals. This has only been examined once to date in subordinate naked mole-rats (oxytocin; Hathaway et al., 2016). Sex steroid hormones (estrogens, androgens) are candidates for driving these differences as they facilitate aggression in many other species (Stetzik et al. 2018; Horton et al. 2014; Soma et al. 2000). Another candidate is dehydroepiandrosterone (DHEA), an androgen precursor produced in the adrenals, implicated in maintaining aggression in reproductively quiescent individuals during non-breeding seasons (Soma et al., 2015). As naked mole-rat subordinates are non-reproductive, this may underly variation in aggression in the absence of high levels of circulating sex hormones. Endocrine profiles can also differentiate breeder from subordinate naked mole-rats (reviewed in Faykoo-Martinez et al., 2021). Plasma estradiol is higher in breeders, whereas RFRP-3 (mammalian ortholog for gonadotropin-inhibitory hormone) is higher in subordinates (Peragine et al., 2017). When a non-reproductive female is removed from the colony, estradiol and progesterone can increase within a week (Faulkes et al., 1990a; Faykoo-Martinez et al., 2018; Swift-Gallant et al., 2015), and continue to increase during the first month (Swift-Gallant et al., 2015), but levels of testosterone do not change (Clarke and Faulkes, 1997; Faulkes et al., 1990a; Faykoo-Martinez et al., 2018; Swift-Gallant et al., 2015). When a non-reproductive male is removed from the colony, testosterone increases within a week, and estradiol and luteinizing hormone increase within a month, but levels of progesterone do not change (Blecher et al., 2020; Faulkes and Abbott, 1991; Faykoo-Martinez et al., 2018; Swift-Gallant et al., 2015). While these studies have helped discern the endocrine signatures of reproductive-status based subordinate and breeder castes, it is not known if the subordinate behavioral phenotypes differ in their hormone levels.

Given the gaps in understanding individual differences within the cooperative naked mole-rat, we investigated behavioral phenotypes in the subordinate caste through classifying subordinates via aggressive and dispersive behaviors rather than the typical working behavior lens. We took a hormonal approach, given that aggressive and dispersive behavior may have endocrine control. We ask two questions: 1) can subordinate naked mole-rats be divided into discrete phenotypes, and 2) do circulating steroid hormone levels underpin these behavioral phenotypes? Since naked mole-rats within a colony tend to be closely related (members of different castes are often full siblings), this is an ideal system to study how and why individuals behave differently within social groups in a controlled, laboratory environment. We tracked the first three litters of six newly-established colonies (N = 178 animals) across three years into adulthood (Figure 1) in captive-living naked mole-rats. To determine behavioral phenotype and variation, we measured behavior towards an opposite-sex conspecific using an outpairing test, a dispersal drive test, and a social dominance test (tube test). To test whether behavioral phenotypes were associated with a distinct hormonal profile, we collected blood samples from a subset of individuals (N = 70) and measured circulating estradiol, progesterone, testosterone, and DHEA levels. Our data were synthesized to determine what milieu of social and biological factors are important for determining the behavioral phenotype profile, which measures are the best predictors, and whether these data are sufficient to categorize behavioral phenotype accurately and discretely.

**Figure 1.**
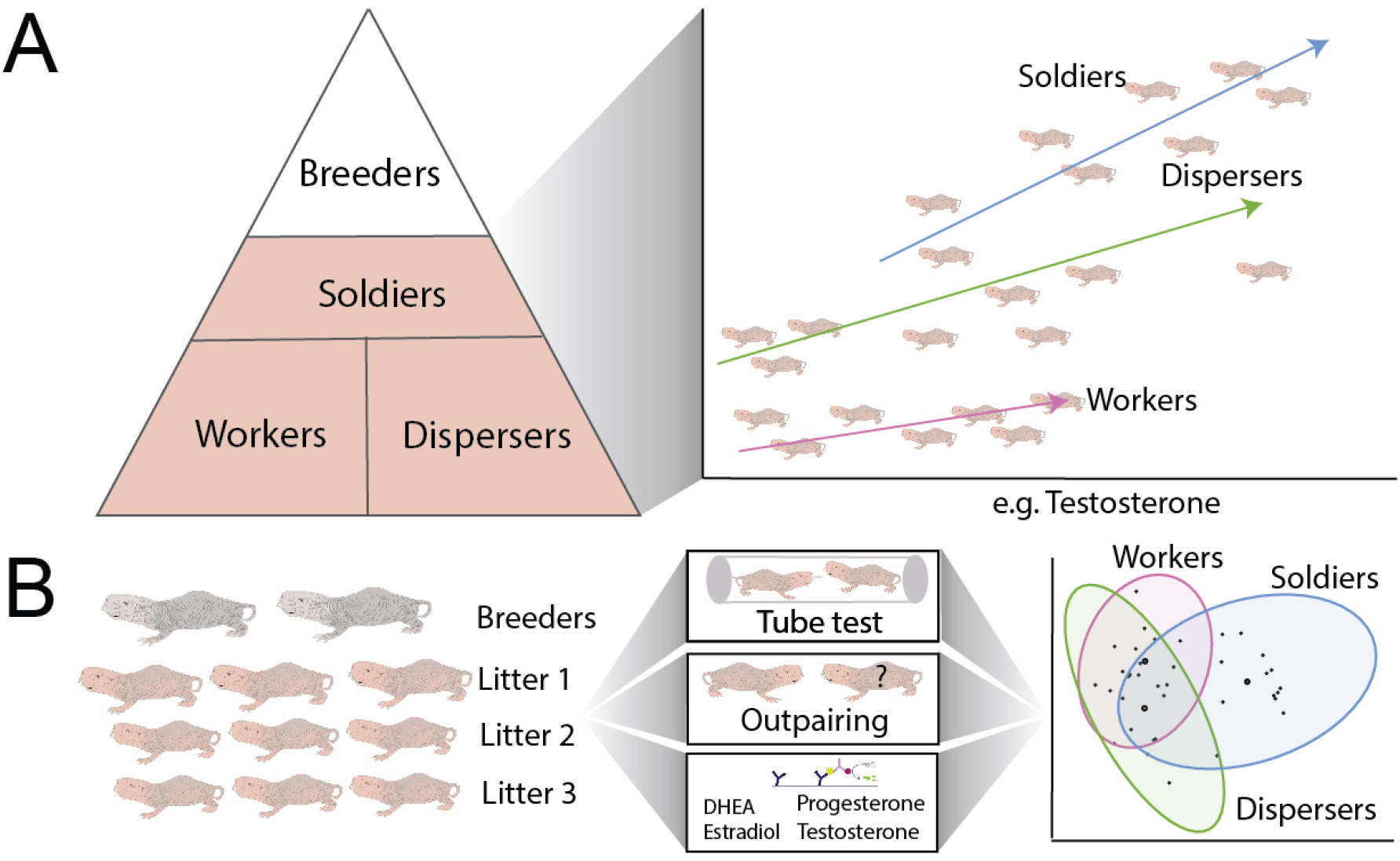
**A)** Naked mole-rat subordinates have been divided into 3 categories: soldiers, workers, and dispersers. It is currently unknown if these categories are continuous or discrete. They are typically defined by biological measures and/or behaviors. **B)** We assessed 6 colonies across the first three litters. We used a social dominance paradigm (tube test), an outpairing test with an unfamiliar opposite-sex conspecific, and measured hormone levels. These data were combined to determine the relationship between hormonal/behavioral milieu across behavioral phenotype and whether these data are sufficient to accurately categorize behavioral phenotype in a discrete manner.

## Methods

### Animals

Six captive colonies (a total of 178 animals) of naked mole-rats maintained in the University of Toronto Mississauga vivarium were used in this study. The colonies were established *de novo* by pairing two opposite-sexed individuals ranging from 12 to 156 months in age. Considering that naked mole-rats reach adulthood within approximately one year and can live for over 37 years, these experimental animals were relatively young adults (Buffenstein and Craft, 2021). All animals were laboratory bred at the University of Toronto Mississauga; the source populations for these animals, and all other laboratory housed research colonies of naked mole-rats, were from field collections in Kenya and Ethiopia >20 years ago (Smith and Buffenstein, 2021). Each colony was housed in polycarbonate caging comprised of a medium (46 cm × 24 cm × 15 cm high) and small (30 cm × 18 cm × 13 cm high) cage connected via polycarbonate tubing, lined with corncob bedding, crinkle paper, and added tubing within the caging. The habitat size was kept constant for the duration of the study. Animal housing rooms were kept between 27°C - 28°C with 50% humidity, and were on a 12-hour light:dark cycle with lights on at 7:00 am. Animals were fed hydrated sweet potato daily and wet Teklad mouse chow three times a week. All work was approved by the University Animal Care Committee (protocol numbers 20011632 and 20011695).

Upon the birth of a litter (average litter size: 10; range: 3-17) each individual pup was assigned a unique color/symbol label for identification using colored permanent markers. Marking occurred at postnatal day 7 to minimize disruption to the litter and cannibalization risk and continued every 3 days until one month of age, when pups were permanently tattooed using an 18 gauge needle and tattoo ink (Ketchum MFG. Co., Cat. No. 392AA). This allowed us to reliably identify each pup and minimize handling associated with re-marking for periods between testing. Pups were tattooed in one area, or a maximum of two areas of the body for larger litters, with each pup in a litter tattooed in a unique location. At six months of age, animals were implanted with a subcutaneous microchip (Avid, Cat. No. 2125, 12 mm). Animals were re-marked with a permanent marker using their unique mark for every testing session to allow identification on video recordings. A maximum of three litters per colony was used for the study (Colony I had only two litters). Once the third litter turned one year old, all three litters went through a battery of behavioral tests to assign behavioral phenotype (described below). All testing took place between 12:00 P.M. and 5:00 P.M. EST.

### Behavior Paradigms

A battery of behavioral paradigms was conducted to categorize the phenotypes of all individuals within each colony. First, all adult animals in the colony were tested using the dispersal test. This testing procedure was adapted from (O’Riain et al., 1996) and is the same as in (Toor et al., 2020). Animals were fed hydrated sweet potato approximately two hours before testing to minimize hunger as motivation to leave the colony. A single hole was opened on the side of the larger cage and a plastic platform (22.86 cm x 30.48 cm) was placed directly under the hole so that animals could easily explore the opening and its surrounding area. When an animal fully exited, its identification was recorded and it was immediately returned to its colony in the cage farthest from the hole. Each trial lasted 30 minutes and three dispersal trials (one per day for three consecutive days) were performed. Animals who exited three or more times during the entire disperser test (combined score across the three trials) were considered dispersers unless they showed aggression during the out-pairing test (see below).

Second, an out-pairing test was administered to all adults (1 year and older) in the colony to determine each individual’s aggressive phenotype. The experimental animal was paired with an unfamiliar opposite-sex animal from a different colony with similar or lower weight. The out-pairing was conducted in a clean 46 cm x 24 cm x 15 cm high cage (medium size) lined with clean bedding. The animals were paired for 10 minutes, and their interactions were recorded using a GoPro Hero 3 camera. In rare cases where the stimulus animal was aggressive to the focal animal, the trial was repeated with a different stimulus animal. A focal animal that was highly aggressive towards the stimulus animal and had to be removed due to risk of injury was labelled a soldier (majority of interactions were stopped before any injury, and no major injuries occurred during this test).

Dispersers were defined as any animal who exited a cumulative of three or more times across three trials of the disperser test and was not aggressive during the out-pairing. Individuals who neither exited the colony during the disperser test nor exhibited aggression during the out-pairing test were considered workers. Workers were not classified based on working type behaviors, but rather the absence of aggression or dispersal-like behavior, because working differentiation between phenotypes remains elusive (Gilbert et al., 2020; Toor et al., 2020).

Third, a social dominance paradigm was conducted as outlined in (Toor et al., 2015) to determine the hierarchical rank of each individual. In the wild, NMRs live in underground chambers connected via tunnels. When two individuals meet in a tunnel, the socially dominant individual will typically walk over the socially submissive individual, which is called a “pass-over”. In the lab, two individuals from the same colony were chosen and simultaneously released at either end of the social dominance apparatus. The animal who passed over the other within the tube was recorded on a matrix. Every animal from a given colony was tested with each and every other animal from its colony in no specific order. A pass-over percentage score was calculated for each animal for each trial where the number of pass-overs performed by a single individual was divided by the total number of possible pairings, multiplied by 100. An estimate of the colony hierarchy could then be constructed by ranking animals within a colony by their pass-over score. In the case of a tie, the order was built by determining which individual within the tied pairs passed over the other.

### Blood Sample Collection

Ideally, four representatives (2 male and 2 female) from each of the 3 phenotypes were collected from each of the six colonies; if an individual of the sex/phenotype combination was unavailable, attempts were made to collect a counterpart from another colony to compensate (Table 1). Representatives were chosen based upon the individuals who displayed the strongest phenotypes (e.g., the most aggressive animals were chosen to represent the soldier phenotypes, the most dispersive for disperser phenotypes, and those with no or very little aggressive/dispersal tendencies as worker phenotypes). In total, 24 soldiers (11 males, 13 females), 25 workers (14 males, 11 females), and 21 dispersers (12 males, nine females) were collected; all collected animals were over one year old and had completed all behavioral testing. Animals were removed from the colony and immediately anesthetized using 4% isoflurane as in Faykoo-Martinez et al. (2018). Animals were then quickly weighed, decapitated, and trunk blood was collected within 5 minutes of removal from the colony. Plasma was stored at -20C until hormone assays were completed.

**Table 1.** Summary of all individuals used for sample collection.

| Colony | Soldiers |  | Dispersers |  | Workers |  |
| --- | --- | --- | --- | --- | --- | --- |
|  | F | M | F | M | F | M |
| L | 3 | 0 | 1 | 3 | 2 | 2 |
| I | 2 | 4 | 3 | 2 | 1 | 4 |
| M | 2 | 2 | 1 | 3 | 3 | 2 |
| F | 1 | 2 | 2 | 2 | 1 | 2 |
| X | 2 | 3 | 1 | 1 | 2 | 2 |
| E | 1 | 0 | 1 | 2 | 2 | 2 |

### Hormone Assays

Estradiol was measured using an ELISA kit (GenWay Biotech, Cat. No. GWB-37E590) with serum diluted 1:5 in buffer. The assay is sensitive to a minimum of 3.94 pg/mL. According to the manufacturer, this ELISA cross-reacts 4.9% with estrone-3-sulfate. The inter-assay coefficient of variation was 5.44% (N=2 plates). Testosterone was measured using an ELISA kit (Enzo Life Sciences, Cat. No. ADI-901-065) with serum diluted 1:10 in buffer. The assay is sensitive to a minimum of 5.67 pg/mL. There is a manufacturer reported assay cross-reactivity with 19-hydroxytestosterone (14.6%), androstenedione (7.30%) and less than 1% of other hormones and metabolites. The inter-assay coefficient of variation was 1.93% (N = 2 plates). Progesterone was measured using an ELISA kit (Cayman Chemical, Cat. No. 582601) with serum diluted 1:10 in buffer. The assay is sensitive to a minimum of 10 pg/mL. There is a manufacturer reported assay cross-reactivity with 17β-estradiol (7.3 %), 5β-pregnan-3α-ol-30-one (6.7 %), pregnenolone (3.5 %), and less than 0.5% with any other hormone or metabolite. The inter-assay coefficient of variation was 17.45% (N = 2 plates). A Synergy-HT-Bio-Tek plate reader was used to measure the ELISA plate absorbance at 405 nm for progesterone and testosterone, and 450 nm for estradiol. All samples were run in duplicate and the average reported. All protocols were followed as per manufacturer specifications.

DHEA was measured using an ^125^I radioimmunoassay kit (MP Biomedicals, Cat. No. 07-230102) which measures both DHEA and DHEA-Sulfate. The assay is sensitive to a minimum of 9 ng/mL. The kit antibody has a cross-reactivity of 100% to DHEA-S and DHEA, 20% to androsterone, 6% to androstenedione, and <1% to estrone, progesterone, testosterone, and 17ßestradiol. This kit requires 25 ul of undiluted plasma per replicate. As such, not all samples had sufficient plasma run in duplicates. The majority were run in duplicate (N = 31 samples) and the others (N = 20) were run as a single replicate. However, RIA replicates tend to be highly stable, and the coefficient of variation between duplicates was 1.68%. As many samples as could be fit on a 100 reaction kit were run. The RIA was run as per manufacturer specifications and samples counted on a Gamma counter.

### Statistical Analysis

First, the associations between behavioral phenotype and within-colony social correlates were analyzed using linear mixed effect models (LMMs). Behavioral phenotype (worker, soldier, disperser) was used as a fixed effect, rank in the hierarchy or litter number were used as the response, and colony was included as a random effect. Then, the effect of behavioral phenotype on each of the four hormones was analyzed using mixed effect models with each hormone as the response variable. We first included additional factors that may be associated with hormone levels. Global models (models with all parameters) therefore included behavioral phenotype, sex, colony mean-adjusted age (months), and colony mean-adjusted weight (g) as fixed effects. Colony was included as a random effect. A single blood sample was taken per animal at sacrifice, not as repeated measures, so animal ID was not included as a random effect. Because some of these fixed effects may be redundant, we calculated the variance inflation factor (VIF) of predictors in the global models for each hormone to test for collinearity between fixed effects. A stringent threshold of a VIF of 3 or higher was set for excluding collinear predictors (Harrison et al., 2018; Zuur et al., 2010). These factors were not collinear in any of the four models. Because we had only four hormones and a low sample size to parameter ratio, P-values were not corrected for false discovery rate (Nakagawa, 2004). Model residuals were normally distributed for concentrations of estradiol (pg/mL) and DHEA (pg/mL), but not for testosterone (pg/mL) nor progesterone (pg/mL). Log transformation ameliorated this issue for the testosterone data but not for the progesterone data. Therefore, estradiol (pg/mL), DHEA (pg/mL) and testosterone (log pg/mL) were analyzed with LMMs while progesterone (pg/mL) was analyzed using a generalized linear mixed effect model (GLMM) with a Gamma family and log link function.

Given that we had several parameters and most of them had no significant effect on hormone levels in the global models, we compared the fit of models with just phenotype as a fixed effect, and models with phenotype and different combinations of the fixed effects (colony always still included as a random effect). Behavioral phenotype was our main factor of interest, so it was always included in the models, and a post-hoc Tukey test was applied to each of the top models (models with the best fit based on corrected Akaike information criterion (AICc) score) to compare the three phenotype groups (worker, soldier, disperser). We compared models using the AICc, which is advantageous for small sample sizes relative to the number of parameters (Burnham et al., 2011) using the package ‘AICcmodavg’ (Mazerolle, 2019).

Hormone levels were then analyzed using outpairing behavior scores. This was to analyze hormones against behavioral measures as a continuum independent of the assigned behavioral phenotypes, to see if there was an association between hormones and behavior outside of the phenotype categories that we assigned. This was done using model selection in the same manner as described above, except behavioral phenotype was replaced with two different continuous measures of behavior. The first was the percent of time spent performing four different behaviors (aggression, sociosexual behavior, prosocial behavior, non-social). Non-social behaviors include, but are not limited to, climbing, digging, self-grooming, in-transit activity, and inactivity. Some outpairing tests were prematurely stopped due to high levels of aggression that could harm the animals. To account for this variation, we used percent of total test time performing a behavior instead of raw duration of behavior. The second measure was the frequency of the four different behaviors, meaning the number of occurrences of a given behavior divided by the total outpairing test time. All behavior measures were converted to z-scores. Because behaviors were our factor of interest, models being compared always had at minimum 1 behavior included (i.e., we are not interested in the sex only, weight only, or age only models and do not compare them). The VIF scores of the factors in each model were checked. In the percent duration behavior models on hormone levels, age and weight were borderline collinear in the DHEA models (VIF of 3.1 and 2.7, respectively) and in the progesterone models (2.7 and 3.2, respectively). No behavioral measures were collinear within any of the behavior percent duration models. In the behavioral frequency models on hormone levels, prosocial and non-social frequency were collinear in all four global models. All factors were kept in the models, but the implications for model selection and interpretation are raised in the results.

Finally, we used a principal component analysis to determine whether our input variables (adjusted age, adjusted weight, hierarchical rank, progesterone, estradiol, testosterone, DHEA, and frequency/percent duration of aggressive, non-social and prosocial sociosexual behaviors) are collectively predictive of behavioral phenotype. The principal components were visualized in 2D and 3D. Principal component scores were further analyzed using a three-way ANOVA for litter, phenotype and sex. P-values were corrected using false discovery rate. Loadings for the input variables for each principal component equal the total sum of squares. Positive valence of the loading indicates a positive relationship between the principal component (and thus any significant variables) and the input variable, and vice versa for negative values. A higher magnitude loading indicates it is contributing greatly to the variation observed in that principal component. This exploratory analysis helps us disentangle which biological measures (i.e. input variables) are influencing our independent variables. For example, soldiers and other subordinates separate on principal component 1. If we detected a significant effect of litter for this principal component, a positive loading for age would indicate that as animals get older, they were more likely to become soldiers and this is ultimately also related to litter number.

Analysis was conducted using R (R Core Team, n.d.). LMMs were constructed using the package ‘nlme’ (Pinheiro et al., 2020) and GLMMs were constructed using the package ‘lme4’ (Bates et al., 2015). Principal component analysis was performed using the base package in R. 3D plots were constructed using ‘rgl’ (Murdoch and Daniel, 2021) and ‘plotly’ (Sievert, 2020). All other figures were constructed using the package ‘ggplot2’ (Wickham, 2016).

## Results

### Behavioral phenotype by social rank and litter

Relative to workers, soldiers were associated with more dominant social rankings within the colony (lower numerical ranking as rank 1 is most dominant; β = -11.88 ± 2.13, t = -5.56, P < 0.001). Dispersers were also associated with more dominant social rankings within the colony compared to workers (β = -6.48 ± 2.15, t = -3.01, P = 0.004). Post hoc tests indicated that dispersers and soldiers also differed (t = 2.45, P = 0.04), with soldiers having more dominant rankings (Figure 2). Relative to workers, soldiers additionally came from earlier litters (β = -1.34 ± 0.16, t = -8.19, P < 0.001). Dispersers were from earlier litters than workers (β = -0.66 ± 0.16, t = -3.98, P < 0.001) but post hoc tests indicated they were also to be from later litters than soldiers (t = 4.05, P < 0.001), hence they are spread across litters but mainly clustered in the second (middle) litter (Figure 2).

**Figure 2.**
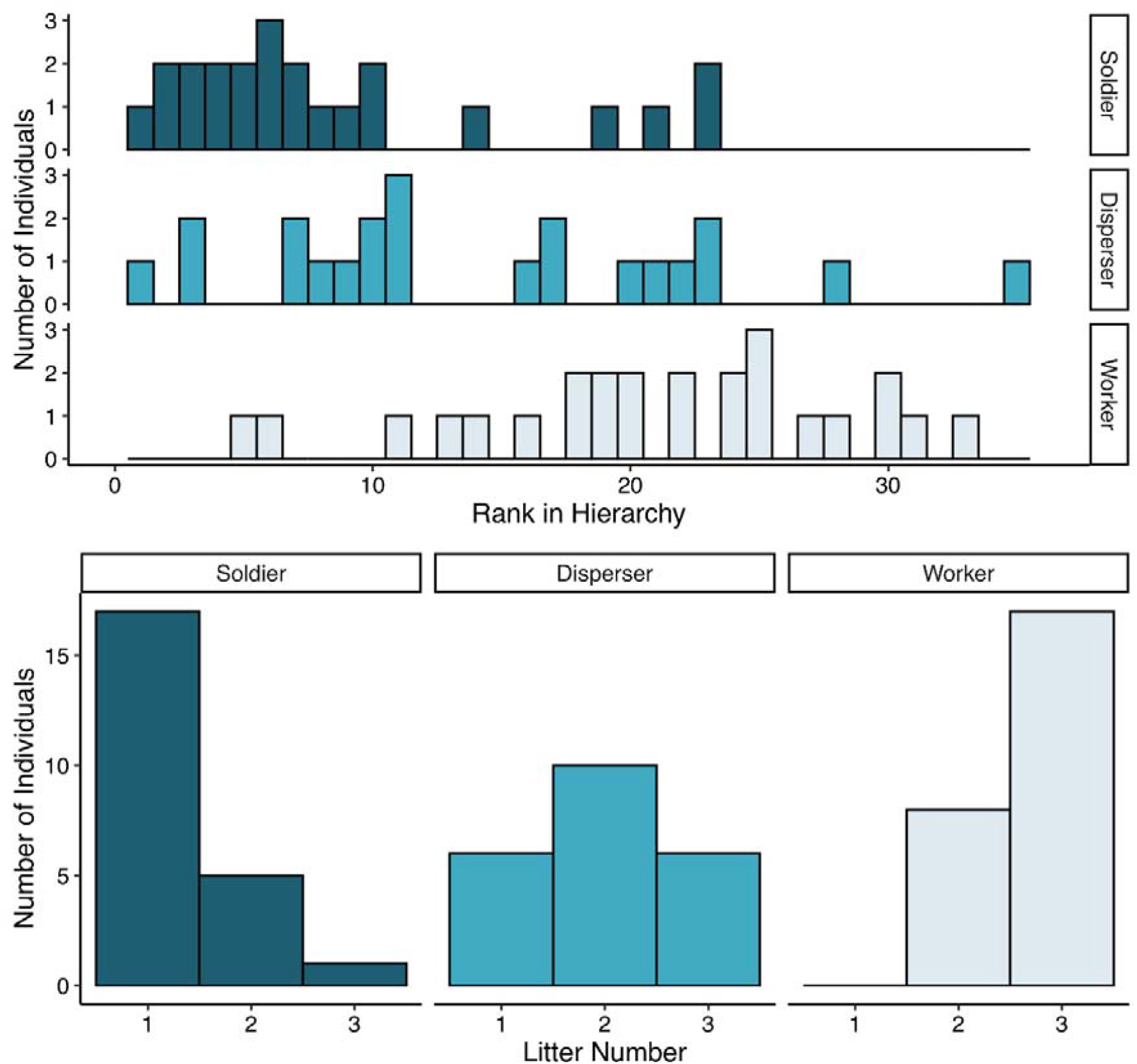
Histograms of number of animals from each phenotype by hierarchical rank and by litter number. The most dominant rank in the hierarchy is 1, with increasing numbers being lower in the rank.

### Hormone levels by behavioral phenotype

We report the results of the top models for each hormone below (Table 2). The results of the global models and model comparisons can be found in Supplementary Tables 1 and 2. For estradiol levels (pg/mL), the top model was the one with behavioral phenotype as the only fixed effect. Phenotype had no detectable effect on estradiol levels (Table 2). The post-hoc Tukey test further showed no difference between any combinations of the three phenotypes: workers and soldiers (t = - 0.53, P = 0.86), workers and dispersers (t = -1.09, P = 0.53), or soldiers and dispersers (t = 0.54, P = 0.85; Figure 3A).

**Table 2.** Results of the top models of the effect of assigned behavioral phenotype on hormone levels. Hormones were estradiol (E), testosterone (T), dehydroepiandrosterone (DHEA), progesterone (P). Fixed effects for behavioral phenotype are dispersers (DIS) and soldiers (SOL) relative to workers as the intercept. Sex is males (M) relative to females as the intercept. Weight is adjusted to the colony mean.

| Response | Fixed effects | Estimate ± SE | t-value | p-value |
| --- | --- | --- | --- | --- |
| E (pg/mL) | PhenotypeDIS | 49.72 ± 45.81 | 1.09 | 0.28 |
| | PhenotypeSOL | 24.25 ± 45.56 | 0.53 | 0.60 |
| T (log pg/mL) | PhenotypeDIS | 0.06 ± 0.10 | 0.54 | 0.59 |
| | PhenotypeSOL | -0.02 ± 0.12 | -0.14 | 0.89 |
|  | **Weight** | **0.82 ± 0.22** | **3.73** | **<0.001** |
| DHEA (pg/mL) | PhenotypeDIS | 611.121 ± 2414.02 | 0.25 | 0.80 |
| | PhenotypeSOL | -3213.58 ± 2727.42 | -1.18 | 0.25 |
|  | **SexM** | **3232.20 ± 1869.17** | **2.17** | **0.04** |
| | Weight | 7406.70 ± 4653.52 | 1.59 | 0.12 |
| P (pg/mL) |  |  | t-value | z-value |
| | PhenotypeDIS | -0.28 ± 0.39 | -0.71 | 0.48 |
|  | **PhenotypeSOL** | **1.42 ± 0.50** | **2.82** | **0.005** |
|  | **SexM** | **-1.28 ± 0.40** | **-3.21** | **0.001** |

**Figure 3.**
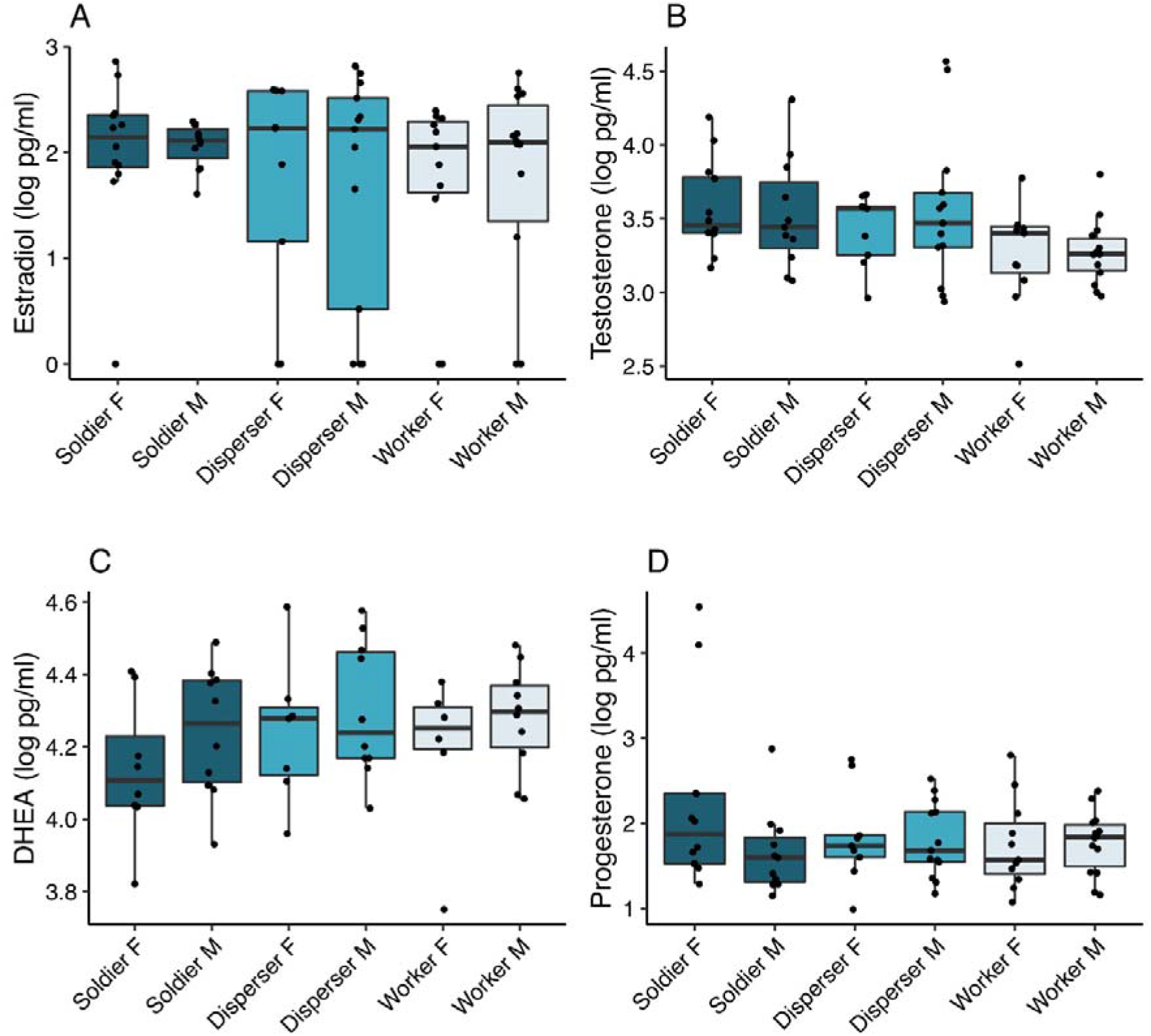
Box plots representing hormone levels (log pg/mL) by behavioral phenotype and sex groups (F = female and M = male). Females had higher progesterone levels than males (P < 0.001) and males had higher DHEA levels than females (P = 0.04). There was no effect of sex on estradiol or testosterone levels. Soldiers had higher progesterone levels than workers (P = 0.02) and dispersers (P = 0.001). No other effects of behavioral phenotype on hormone levels were detected.

For testosterone levels (log pg/mL), the top model was the one with phenotype and weight as fixed effects. Weight had a positive effect on testosterone, with heavier animals having higher testosterone levels (P < 0.001). When accounting for the effect of weight, behavioral phenotype had no detectable effect on testosterone levels (Table 2). The post-hoc contrasts additionally showed no difference between workers and soldiers (t = 0.14, P = 0.99), workers and dispersers (t = -0.54, P = 0.85), or soldiers and dispersers (t = 0.74, P = 0.74; Figure 3B).

For DHEA levels (pg/mL), the top models were the ones with phenotype and sex as fixed effects and phenotype, sex, and weight as fixed effects, which did not meaningfully differ (βAICc = 0.01, Supplementary Table 2). Sex had an effect on DHEA levels, with males having higher levels than females (P = 0.04; Table 2). Weight and behavioral phenotype had no detectable effect on DHEA levels (Table 2). Post-hoc contrasts showed no difference between workers and soldiers (t = 1.18, P = 0.47), workers and dispersers (t = -0.25, P = 0.97), or soldiers and dispersers (t = 1.73, P = 0.21; Figure 3C).

For progesterone levels (pg/mL), the top model was the one with phenotype and sex. Males had markedly lower progesterone levels than females (t = -3.32, P = 0.001). Further, there was an effect of behavioral phenotype, and post-hoc tests indicated that soldiers had higher progesterone levels than workers (z = -2.82, P = 0.01) and dispersers (z = -3.60, P = 0.001). Workers and dispersers did not differ from each other (z = 0.71, P = 0.76; Figure 3D).

### Hormone levels by behavior as a continuum

We report the results of the top models for each hormone below. The results of the global models and model comparisons can be found in Supplementary Tables 3 and 4. For estradiol levels (pg/mL), the top model for behavior percent duration was the prosocial duration model, and the top model for behavior frequency was the aggression frequency model. In the estradiol and behavior duration models, the prosocial model did not meaningfully differ from the subsequent two top models, which were the aggression and prosocial + sociosexual models (ΔAICc < 2, Supplementary Table 4). Prosocial behavior had no detectable relationship with estradiol levels (Table 3). In the estradiol and behavior frequency models, the top six models all included aggression, and once aggression was dropped in the seventh-ranked model, the AICc score increased slightly (ΔAICc = 2.38). However, aggression had only a marginal positive association with estradiol levels (P = 0.08; Table 3).

**Table 3.**
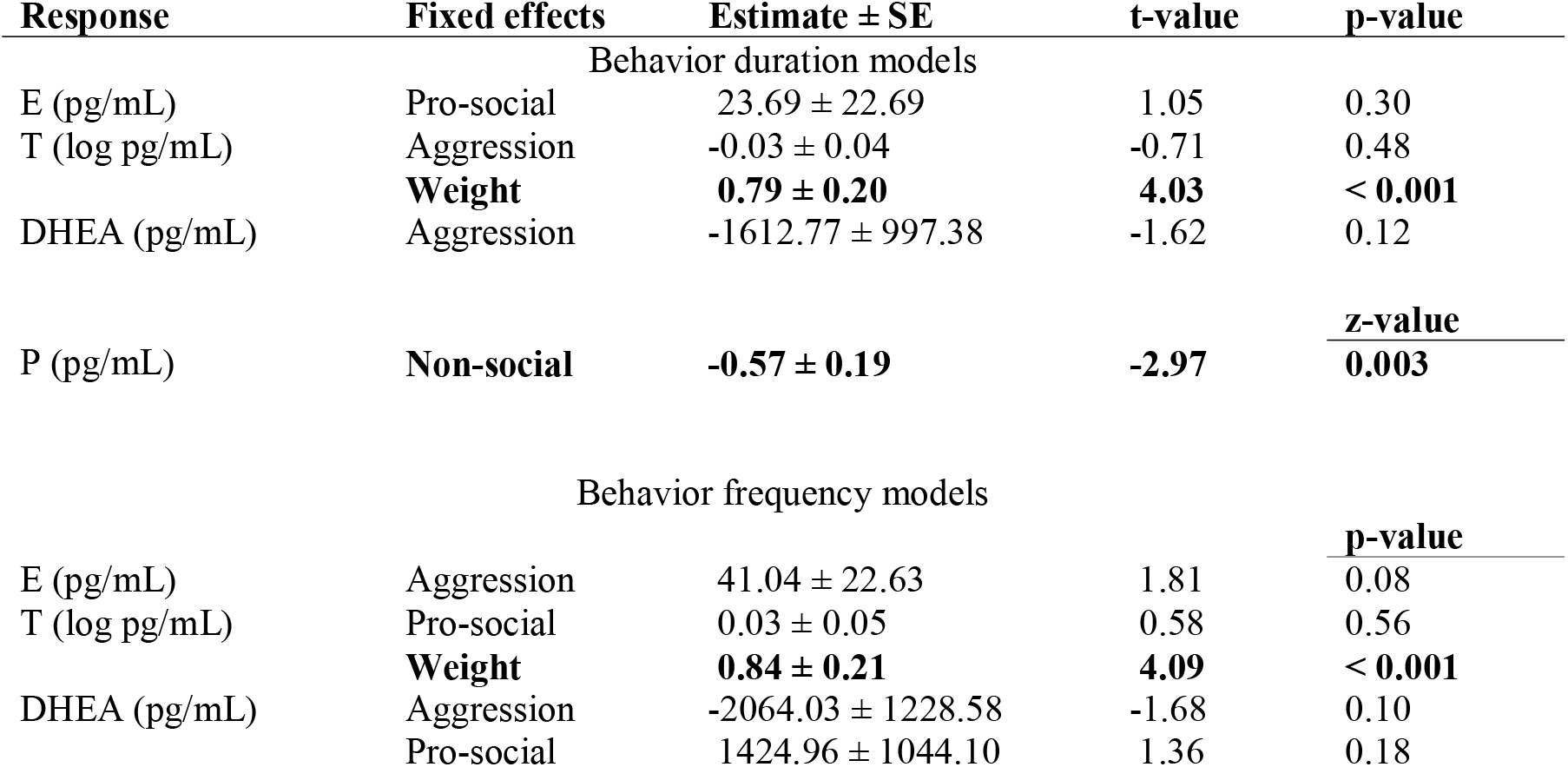
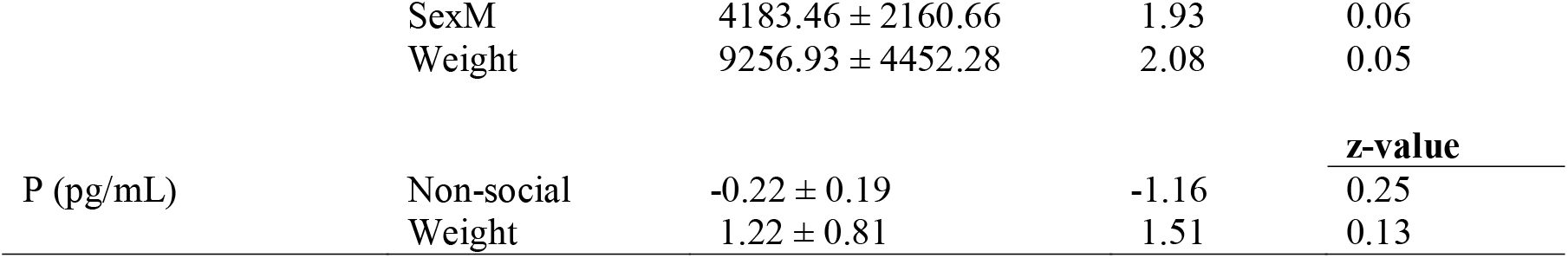
Results of the top models of the effect of behavior scores on hormone levels. Hormones were estradiol (E), testosterone (T), dehydroepiandrosterone (DHEA), progesterone (P). The behavior durations models are based on the z-scored percent duration of total outpairing test time, and the behavior frequencies models are based on the z-scored occurrences divided by the total outpairing test time. Sex is males (M) relative to females as the intercept. Weight is adjusted to the colony mean.

**Table 4.** Loadings of input variables for the top three principal components. Loadings are rounded to 3 significant digits. Loadings for all principal components can be accessed in *Supplementary File 1. PC=Principal component*

| **Input variable** | **PC1** | **PC2** | **PC3** |
| --- | --- | --- | --- |
| Colony-centered adjusted weight | 0.23 | -0.45 | 0.14 |
| Colony-centered adjusted age | 0.13 | -0.39 | -0.15 |
| Hierarchical rank | 0.28 | -0.36 | 0.05 |
| log Progesterone (pg/mL) | -0.32 | -0.23 | 0.25 |
| log Estradiol (pg/mL) | -0.35 | -0.231 | -0.21 |
| log Testosterone (pg/mL) | -0.40 | -0.20 | -0.11 |
| log DHEA (pg/mL) | -0.37 | -0.19 | -0.16 |
| Aggression Percent | -0.01 | -0.19 | -0.27 |
| Aggression Frequency | 0.05 | -0.17 | -0.40 |
| Neutral Percent | -0.35 | -0.14 | 0.16 |
| Neutral Frequency | -0.38 | 0.01 | -0.04 |
| Pro-social Percent | 0.07 | 0.06 | -0.57 |
| Pro-social Frequency | 0.09 | 0.09 | -0.40 |
| Sociosexual Percent | -0.02 | 0.18 | -0.23 |
| Sociosexual Frequency | -0.23 | 0.45 | -0.01 |

For testosterone levels (log pg/mL), the top model for behavior percent duration was the aggression duration and weight model. Aggression had no detectable relationship with testosterone levels (P = 0.48); however, weight was again positively associated with testosterone levels (P < 0.001; Table 3). For testosterone levels and behavior frequency, the top model was the prosocial frequency and weight model, but prosocial frequency had no association with testosterone levels (P = 0.56). Weight again had a positive association with testosterone levels (P < 0.001; Table 3).

With regard to DHEA levels (pg/mL), the top model for behavior percent duration was the aggression duration model, but aggression duration was not associated with DHEA levels (P = 0.12; Table 3). For DHEA and behavior frequency, the top model was the aggression, prosocial, sex, and weight model. Behavior frequency did not have an association with DHEA levels, and sex and weight had only a weak marginal relationship with DHEA levels (P = 0.06 and P = 0.05 respectively; Table 3).

For progesterone levels (pg/mL), the top model for behavior percent duration was the non-social duration model. Non-social percent duration had a negative association with progesterone levels (P = 0.003; Table 3), indicating animals with higher progesterone spent less time being non-social. For progesterone and behavior frequency, the top model was the non-social frequency and weight model, though this model did not meaningfully differ from the next six top models (all ΔAICc < 2, Supplementary Table 4). Non-social frequency had no detectable association with progesterone levels (P = 0.25), nor did weight (P = 0.13; Table 3).

### Predicting behavioral phenotype by principal component analysis

A principal component analysis revealed a distinct separation of soldiers from dispersers and workers on the first principal component 1 (33.3% of variation explained), whereas dispersers and workers begin to separate on the second principal component (17.6% of variation explained) (Figure 4A). While the clustering of soldiers revealed a clearer phenotype separation, differences between dispersers and workers do emerge. We also visualized the first three principal components (third principal component, 9.93% of variation explained) in a 3D principal component analysis plot to further explore the variation between phenotypes (Supplementary Figure 1). A 3-way ANOVA of the sex, litter and phenotype effects on PC scores revealed significant effects in principal components 1 and 2, but not 3.

**Figure 4.**
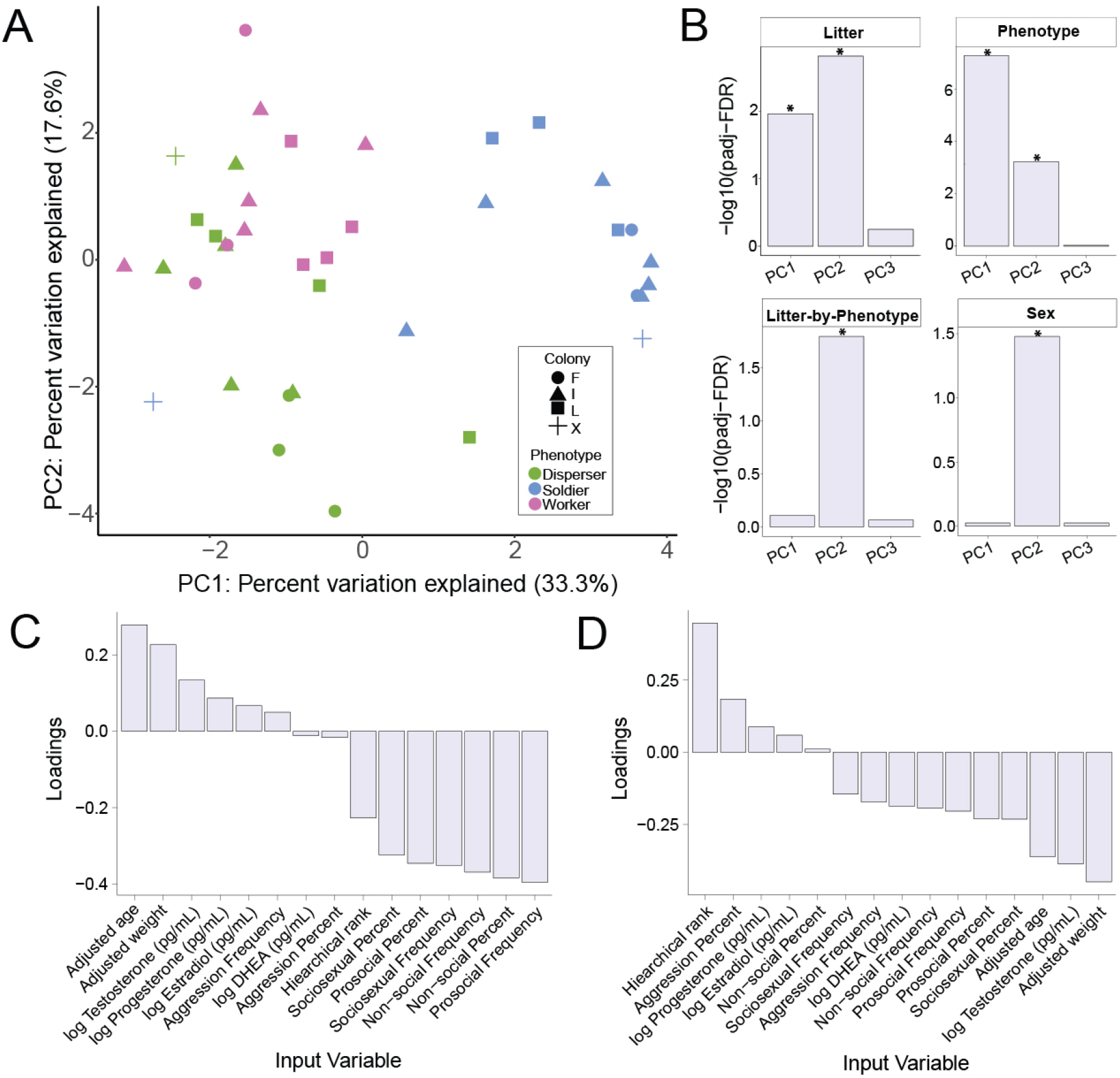
**A)** A principal component analysis of dependent variables measured (age, weight, hormones and behavior). Colony is indicated by shape and phenotype is indicated by colour. **B)** Bar plots representing the -log10(p-adjusted) using a false detection rate correction the 3-way ANOVA of litter, phenotype and sex on principal component scores. A significant effect of litter (padj=0.01) and phenotype (padj<0.001) was observed for principal component 1. A significant effect of litter (padj=0.001), phenotype (padj<0.001), sex (padj=0.03) and a litter-by-phenotype interaction (padj=0.02) was observed for principal component 2. No significant effects were observed for principal component 3. **C)** Loadings for the first principal component rank-ordered by value. **D)** Loadings for the second principal component rank-ordered by value. *PC=principal component, padj-FDR=False discovery rate adjusted p-value*

The main effect of litter (padj=0.01) and phenotype (padj<0.001) in principal component 1 can be primarily attributed to a negative association with pro-social frequency (−0.39), non-social percent duration (−0.38), non-social frequency (−0.37), sociosexual frequency (−0.35), pro-social percent (−0.35) and sociosexual percent duration (−0.32) (Figure 4C). The main effect of litter (padj=0.001), phenotype (padj<0.001), sex (padj=0.03) and a litter-by-phenotype interaction (padj=0.020) in principal component 2 can be primarily attributed to associations with adjusted weight (−0.46), testosterone (−0.39), adjusted age (−0.36) and hierarchical rank (0.45). Loadings for the top three principal components are summarized in Table 4. All ANOVA outputs and principal component loadings can be found in Supplementary File Figure 5.

## Discussion

We had two objectives in our study: first, to determine whether workers, soldiers, and dispersers are distinct phenotypes within a naked mole-rat colony or are simply on a behavioral spectrum; and second, to investigate the social and hormonal correlates of this behavioral variation. Our data support predictions that naked mole-rat subordinate behavioral phenotypes tend to be distinct based on social correlates, albeit still existing on a large spectrum. Furthermore, steroid hormones are less important in differentiating between the behavioral phenotypes. Direct comparisons of the assigned behavioral phenotypes revealed social differences but few hormonal differences. The three behavioral phenotypes significantly differed in hierarchical rank within their respective colony, where soldiers were the most dominant of the group, and workers the least. Soldiers were from earlier litters and workers from later litters, whereas dispersers were born across all three litters. Overall, behavioral phenotypes were not associated with a distinct hormonal profile. No hormonal differences between social phenotypes were found with respect to estradiol, testosterone, or DHEA levels. Soldiers had higher progesterone levels; however, this effect appeared to be driven by a few particularly high individuals, and many soldiers had progesterone levels that were equally low or lower than individuals from other groups.

Upon close inspection of various naked mole-rat characteristics, including hormones, behavior, and physical traits via principal component analysis, soldiers seemed to differentiate themselves from workers and dispersers, although overlap is evident. Variables that contributed to differences between soldiers and workers include weight, testosterone, prosocial behavior, and aggression, albeit loosely associated. Variables that contributed to differences between workers and dispersers include testosterone, aggression, weight, and hierarchical rank. Data from the general linear models and principal component analysis suggest that the relationship between behavioral phenotype and testosterone is driven by the correlation between testosterone and body weight, which is associated with phenotype. However, the directionality of testosterone and body composition in naked mole-rats is not clear cut. Environmental pressures can alter the interaction between testosterone and body weight. For example, arctic ground squirrels survive hibernation in the tundra with higher endogenous levels of androgen receptor relative to other ground squirrels, allowing them to increase their muscle mass prior to winter (Boonstra et al., 2014). When accounting for the effect of body weight, there is no relationship between phenotype and testosterone level (Table 2).

Our finding that behavioral phenotype, rank, weight, and litter number are all related is consistent with previous work. Heavier and older mole-rats tend to be more highly ranked in the hierarchy than smaller and younger mole-rats, specifically when using the social dominance paradigm (Toor et al., 2015). This association is consistent with observations of undisrupted colony behaviors where aggressive behavior and higher body weight of individuals is associated with higher positions within a dominance hierarchy in both males and females (Clarke and Faulkes, 1997, 1998). Additionally, the first litter in a naked mole-rat colony typically grows faster than subsequent litters, most likely to quickly join in colony maintenance until their younger siblings take over maintenance roles (Bennett et al., 1991; Jarvis, 1991). Thus, it is not surprising that we found a strong association with the larger and more aggressive soldier phenotype naked mole-rats to be mainly from the first litter and higher in the colony hierarchy. Furthermore, there is evidence of dispersing naked mole-rats being generally larger or heavier than their colony mates (O’Riain et al., 1996; Braude, 2000), albeit with some mixed results with female disperser-types being heavier than worker-types, but not for males (Toor et al., 2020). Given the previously established relationship between body mass and hierarchy, it is interesting that we found dispersers to be spread across the litters and the hierarchy. We acknowledge that there is a certain circularity found within this paper: aggression (presence/absence, duration) is essential in defining these social phenotypes. Thus, it is hardly surprising we found significant differences in aggression between soldiers and non-soldiers. However, these findings still play a crucial role in the study of naked mole-rats by offering an example of stability in traits through a combination of outpairing aggression and in-colony social dominance.

Progesterone was the only hormone found to be higher in soldiers than in workers and dispersers, and no other hormonal differences were found among phenotypes. While multiple studies have looked at hormonal differences between reproductive and non-reproductive naked mole-rats, hormone differences within the non-reproductive caste have not been extensively explored. Generally, subordinate naked mole-rats have lower testosterone, estradiol, and progesterone levels than breeders (Clarke and Faulkes, 1997; Faulkes et al., 1990b; Faykoo-Martinez et al., 2018; Swift-Gallant et al., 2015). Urinary testosterone is strongly negatively correlated with dominance rank in both sexes, and subordinate females with the highest testosterone levels became reproductively active upon queen removal (Clarke and Faulkes, 1997). Large and dominant non-reproductive females additionally exhibit increased progesterone levels after queen removal (Clarke and Faulkes, 1997). Increases in progesterone are also found when subordinate females are removed from the colony and singly housed, and levels continue to increase once they are paired with a male mate (Blecher et al., 2020; Swift-Gallant et al., 2015). Our study finding soldiers to have higher progesterone levels may indicate dominant individuals within a colony are primed and ready to take over as breeders, although it is unexpected that no explicit differences in testosterone were found between the three phenotypes; rather, testosterone differences were more directly associated with weight. Given that aggression is such an important part of deciphering the three phenotypes, we quantified levels of DHEA, which is linked to aggression and territorial defense in non-breeding individuals in other species (Boonstra et al., 2008; Rendon and Demas, 2016). DHEA is released from the adrenals and converted to androstenedione in the brain, which can then be converted to testosterone or directly aromatized into estradiol (reviewed Soma et al., 2015). Since subordinate naked mole-rats are essentially permanently non-breeding, in the absence of changes in testosterone or estradiol, we predicted that DHEA may facilitate differences in aggression among the behavioral phenotypes. However, this did not appear to be the case, as DHEA did not differ among phenotypes (Table 2; Figure 3), nor did DHEA have any association with aggression levels (Table 3). We did not test another critical adrenal hormone, cortisol, but our previous work has shown that there are no rank-related differences in fecal cortisol metabolites in this species (Edwards et al., 2020). The fact that rank is strongly tied to behavioral phenotype makes it unlikely that cortisol would separate out by phenotype either.

Since there was no strong endocrine profile associated with behavioral phenotype or continuous measures of behavior in naked mole-rats, this variation is most likely facilitated by another biological mechanism. For example, there could be behavioral phenotype-related differences in brain receptor expression, or in conversion rate of DHEA or testosterone to estradiol (e.g. differences in aromatase). It is also possible that these differences in aggression and other aspects of social behavior in subordinate naked mole-rats are driven by other neuroendocrine mechanisms beyond steroid hormones. Workers have been found to have more oxytocin neural activity in the paraventricular nucleus of the hypothalamus than do soldiers (Hathaway et al., 2016). Because only the present study and Hathaway et al. 2016 have probed the physiology of subordinate behavioral phenotypes in naked mole-rats, there are many untested possibilities of mechanisms, neuroendocrine or otherwise, that could be driving this behavioral variation.

The principal component analysis gave insight into the variables that separate the three phenotypes from one another but there was overlap, especially when comparing workers and dispersers (see Toor et al., 2020 for details on variation in dispersal tendencies). Many individuals fall in between the phenotypes during the characterization stage. For example, during the disperser test, some naked mole-rats would show mild exploratory behaviour by only exiting the colony once or twice. Others would be highly explorative and exit more than three times, but were incredibly aggressive during outpairing. During outpairing, some would even show sporadic aggression and ample prosocial behaviour towards the unfamiliar conspecific but others would largely ignore it instead. Indeed, out of a total of 178 animals, we aimed to recruit four of the best qualifiers of each social phenotype (2 male, 2 female) from each colony: the most aggressive soldiers, the most explorative dispersers, and the most unaggressive and unexplorative workers. However, it is clear when looking at the recruitment numbers (Table 1) that an even spread of “the best” was not necessarily even, and the definition of “the best” would vary from colony to colony. Thus, although our 70 qualifying naked mole-rats are in as discrete categories as we could get there is still considerable overlap. The overlap rings true for disperser and workers moreso than the soldiers, who have a more discrete categorization. This variation qualitatively indicates that the phenotypes are more on a spectrum rather than in distinct classes. There is evidence of some behaviors falling on a spectrum and others being bimodal, but when colony dynamics change by removing individuals, individuals appear to change their behavior to compensate for the missing colony members (Mooney et al., 2015). Colony members who fall in between two phenotypes (e.g. soldier-like dispersers, or disperser-like workers) may be the ones who transition into various roles depending on the needs of the colony, but further research will need to be conducted to confirm this. Interestingly, sociosexual duration and adjusted weight differentiate dispersers from workers, which corroborates past literature on dispersers being more sexually interested in opposite-sexed conspecifics and generally being larger than workers (O’Riain et al., 1996; Toor et al., 2020). Dispersers and disperser-like workers could be more opportunistically sociosexual outside of the colony, which their name implies, where they may have more opportunities to mate away from the large, high ranking soldiers who are more likely to become reproductive following the death of the breeders (Clarke and Faulkes, 1997). However, it is important to note that disperser females show more agonistic, breeder-like behaviors in the colony than female workers (Toor et al., 2020). Further research could clarify if this is yet another example of the behavioral spectrum exhibited by naked mole-rats or a unique characteristic of the disperser phenotype.

The present work identifies behavioral phenotype differences within the non-reproductive caste in naked mole-rats and measures that help distinguish soldiers, dispersers, and workers. Although few hormonal differences were found, other biological measures at play likely underlie the behavioral variation in this unique mammalian eusocial system. Given that we already know oxytocin-expressing cells differ between soldiers and workers, it is possible that other neural enzyme/receptor expression differences are at play between phenotypes or that neuropeptides, not steroid hormones, drive these behavioral differences. Based on the metrics in this study, soldiers appear to form a discrete category, but workers and dispersers have more overlap with each other. Further, within each category, there is individual variation. By continuing to investigate measures beyond body composition and hormones, we can further establish the biological milieu that individuates subordinate behavioral phenotypes and, potentially, find a path towards characterizing these phenotypes while minimizing within-group variation.

## Funding Sources

This work was supported by an Ontario Graduate Scholarship and a Natural Sciences and Engineering Research Council of Canada (NSERC) Postgraduate Scholarship - Doctoral (PGS D) grant to IT, NSERC CGS-D, OGS and ACM/Intel Computational Fellowship to MFM, a University of Toronto Mississauga Postdoctoral Award to PDE, NSERC grant (RGPIN-2016-05540) to RB, and NSERC grants (RGPIN 2018-04780 and RGPAS 2018-522465), CIHR grant (02003PJT-437197) and an Ontario Early Researcher Award to MMH.

## Notes

### Competing Interest Statement

The authors have declared no competing interest.

