## Supplementary Tables for "Hormones do not make the mole-rat: no steroid hormone signatures of subordinate behavioral phenotypes"

Table A1. Results of the global models. Fixed effects for behavioral phenotype are dispersers (DIS) and soldiers (SOL) relative to workers as the intercept. Sex is males (M) relative to females as the intercept. Age and weight are adjusted to the colony mean.

| **Response** | **Fixed effects** | **Estimate ± SE** | **t-value** | **p-value** |
| --- | --- | --- | --- | --- |
| Estradiol | PhenotypeDIS | 47.76 ± 52.94 | 0.90 | 0.37 |
|  | PhenotypeSOL | 19.69 ± 64.72 | 0.30 | 0.76 |
|  | SexM | 11.00 ± 38.84 | 0.28 | 0.78 |
|  | Age | 88.35 ± 167.04 | 0.53 | 0.78 |
|  | Weight | -53.26 ± 133.75 | -0.40 | 0.60 |
| Testosterone | PhenotypeDIS | 0.04 ± 0.11 | 0.46 | 0.65 |
|  | PhenotypeSOL | -0.03 ± 0.13 | -0.22 | 0.83 |
|  | SexM | 0.04 ± 0.08 | 0.52 | 0.60 |
|  | Age | 0.11 ± 0.28 | 0.37 | 0.71 |
|  | **Weight** | **0.77 ± 0.26** | **3.00** | **0.004** |
| DHEA | PhenotypeDIS | 873.84 ± 2457.68 | 0.36 | 0.72 |
|  | PhenotypeSOL | -2604.86 ± 2914.40 | -0.89 | 0.38 |
|  | **SexM** | **4038.04 ± 1850.16** | **2.18** | **0.03** |
|  | Age | -4848.26 ± 8648.63 | -0.56 | 0.58 |
|  | Weight | 9831.26 ± 6359.51 | 1.55 | 0.13 |
|  |  | **Estimate ± SE** | **t-value** | **z-value** |
| Progesterone | PhenotypeDIS | -0.28 ± 0.46 | -0.60 | 0.55 |
|  | **PhenotypeSOL** | **1.48 ± 0.70** | **2.12** | **0.04** |
|  | **SexM** | **-1.28 ± 0.40** | **-3.18** | **<0.001** |
|  | Age | -1.34 ± 1.81 | -0.74 | 0.46 |
|  | Weight | 0.71 ± 1.23 | 0.57 | 0.57 |

Table A2. Model comparisons with each hormone as the response variable. Age and weight are adjusted to the colony mean.

| **Response** | **Fixed effects** | **K** | **AICc** | **ΔAICc** |
| --- | --- | --- | --- | --- |
| Estradiol | **Phenotype** | **5** | **920.18** | **0.00** |
| (pg/mL) | Phenotype+age | 6 | 922.50 | 2.32 |
|  | Phenotype+sex | 6 | 922.51 | 2.33 |
|  | Phenotype+weight | 6 | 922.56 | 2.38 |
|  | Phenotype+sex+age | 7 | 924.90 | 4.72 |
|  | Phenotype+sex+weight | 7 | 924.97 | 4.80 |
|  | Phenotype+sex+age+weight | 8 | 927.33 | 7.16 |
| Testosterone | **Phenotype+weight** | **6** | **46.45** | **0.00** |
| (log pg/mL) | Phenotype+sex+weight | 7 | 48.62 | 2.17 |
|  | Phenotype+sex+age+weight | 8 | 51.03 | 4.57 |
|  | Phenotype+age | 6 | 55.38 | 8.92 |
|  | Phenotype | 5 | 57.46 | 11.00 |
|  | Phenotype+sex+age | 7 | 57.70 | 11.24 |
|  | Phenotype+sex | 6 | 59.73 | 13.27 |
| DHEA | **Phenotype+sex** | **6** | **1056.92** | **0.00** |
| (pg/mL) | Phenotype+sex+weight | 7 | 1056.93 | 0.01 |
|  | Phenotype | 5 | 1058.29 | 1.37 |
|  | Phenotype+weight | 6 | 1058.94 | 2.02 |
|  | Phenotype+sex+age | 7 | 1058.97 | 2.04 |
|  | Phenotype+sex+age+weight | 8 | 1059.51 | 2.59 |
|  | Phenotype+age | 6 | 1060.31 | 3.38 |
| Progesterone | **Phenotype+sex** | **6** | **889.44** | **0.00** |
| (pg/mL) | Phenotype+sex+age | 7 | 891.67 | 2.23 |
|  | Phenotype+sex+weight | 7 | 891.87 | 2.43 |
|  | Phenotype+sex+weight+age | 8 | 893.89 | 4.45 |
|  | Phenotype | 5 | 897.28 | 7.85 |
|  | Phenotype+age | 6 | 899.45 | 10.02 |
|  | Phenotype+weight | 6 | 899.58 | 10.14 |

Table A3. Results of the global models for behavior durations, measured in the z-scored percent duration of total outpairing test time, and results of the global models for behavior frequencies, measured in the z-scored occurrences divided by the total outpairing test time. Age and weight are adjusted to the colony mean.

| **Response** | **Fixed effects** | **Estimate ± SE** | **t-value** | **p-value** |
| --- | --- | --- | --- | --- |
|  | Behavior duration models | | | |
| Estradiol | Aggression | 20.98 ± 25.14 | 0.83 | 0.55 |
| (pg/mL) | Non-social | -12.93 ± 30.76 | -0.42 | 0.41 |
|  | Pro-social | 38.54 ± 28.74 | 1.34 | 0.19 |
|  | Sociosexual | -14.55 ± 31.57 | -0.46 | 0.65 |
|  | SexM | 44.36 ± 47.15 | 0.94 | 0.35 |
|  | Age | 83.35 ± 194.66 | 0.42 | 0.67 |
|  | Weight | -52.27 ± 151.14 | -0.38 | 0.71 |
| Testosterone | Aggression | -0.02 ± 0.04 | -0.50 | -0.62 |
| (log pg/mL) | Non-social | 0.02 ± 0.06 | 0.40 | 0.69 |
|  | Pro-social | 0.03 ± 0.06 | 0.45 | 0.66 |
|  | Sociosexual | -0.02 ± 0.06 | 0.42 | 0.67 |
|  | SexM | 0.11 ± 0.09 | 1.21 | 0.23 |
|  | Age | 0.02 ± 0.33 | 0.07 | 0.95 |
|  | **Weight** | **0.83 ± 0.29** | **2.91** | **0.006** |
| DHEA | Aggression | -953.81 ± 1076.40 | -0.89 | 0.38 |
| (pg/mL) | Non-social | 1410.89 ± 1634.29 | 0.86 | 0.40 |
|  | Pro-social | -1326.09 ± 1470.07 | -0.90 | 0.38 |
|  | Sociosexual | 431.93 ± 1991.47 | 0.22 | 0.83 |
|  | SexM | 3202.31 ± 2369.89 | 1.60 | 0.12 |
|  | Age | -11723 ± 9892.51 | -1.19 | 0.25 |
|  | Weight | 11452.08 ± 7235.06 | 1.99 | 0.06 |
|  |  |  |  | **z-value** |
| Progesterone | Aggression | -0.18 ± 0.16 | -1.14 | 2.55 |
|  | Non-social | -0.46 ± 0.28 | -1.69 | 0.09 |
|  | Pro-social | -0.01 ± 0.19 | -0.06 | 0.95 |
|  | Sociosexual | -0.24 ± 0.24 | -0.98 | 0.33 |
|  | SexM | -0.55 ± 0.38 | -1.45 | 0.14 |
|  | Age | -0.09 ± 1.59 | -0.06 | 0.95 |
|  | Weight | 0.19 ± 1.26 | 0.15 | 0.88 |
|  | Behavior frequency models | | | |
|  |  |  |  | **p-value** |
| Estradiol | Aggression | 41.34 ± 25.98 | 1.59 | 0.12 |
| (pg/mL) | Non-social | 36.89 ± 54.64 | 0.68 | 0.50 |
|  | Pro-social | -18.05 ± 61.10 | -0.30 | 0.77 |
|  | Sociosexual | -4.36 ± 36.99 | -0.12 | 0.91 |
|  | SexM | 51.26 ± 46.60 | 1.10 | 0.28 |
|  | Age | -17.49 ± 211.14 | 0.08 | 0.93 |
|  | Weight | -44.97 ± 148.63 | -0.30 | 0.76 |
| Testosterone | Aggression | 0.007 ± 0.05 | 0.14 | 0.89 |
| (log pg/mL) | Non-social | -0.09 ± 0.11 | 0.86 | 0.40 |
|  | Pro-social | 0.14 ± 0.12 | 1.17 | 0.24 |
|  | Sociosexual | -0.03 ± 0.06 | -0.45 | 0.65 |
|  | SexM | 0.13 ± 0.09 | 1.44 | 0.16 |
|  | Age | -0.11 ± 0.37 | -0.29 | 0.77 |
|  | **Weight** | **0.91 ± 0.28** | **-3.30** | **0.002** |
| DHEA | Aggression | -2308.15 ± 1415.83 | -1.63 | 0.11 |
|  | Non-social | 2588.05 ± 2483.69 | 1.04 | 0.31 |
|  | Pro-social | -685.26 ± 2838.32 | -0.24 | 0.81 |
|  | Sociosexual | -1132.23 ± 1899.17 | -0.60 | 0.56 |
|  | **SexM** | **5358.81 ± 2431.56** | **2.20** | **0.04** |
|  | Age | -8623.09 ± 10059.05 | -0.86 | 0.40 |
|  | Weight | 13677.46 ± 6972.32 | 1.96 | 0.06 |
|  |  |  |  | **z-value** |
| Progesterone | Aggression | -0.09 ± 0.23 | -0.39 | 0.69 |
|  | Non-social | 0.16 ± 0.55 | 0.30 | 0.76 |
|  | Pro-social | -0.40 ± 0.57 | -0.70 | 0.49 |
|  | Sociosexual | -0.14 ± 0.28 | -0.50 | 0.62 |
|  | SexM | -0.46 ± 0.37 | -1.22 | 0.22 |
|  | Age | 0.71 ± 1.78 | 0.40 | 0.69 |
|  | Weight | 0.61 ± 1.29 | 0.47 | 0.64 |

Table A4. Model comparisons with each hormone as the response variable. The top 7 models that include any behavioral measures are reported. Age and weight are adjusted to the colony mean.

| **Response** | **Fixed effects** | **K** | **AICc** | **ΔAICc** |
| --- | --- | --- | --- | --- |
|  | Behavior duration models | | | |
| Estradiol | **Pro-social** | **4** | **740.31** | **0.00** |
| (pg/mL) | Aggression | 4 | 742.07 | 1.76 |
|  | Pro-social+sociosexual | 5 | 742.28 | 1.97 |
|  | Sociosexual | 4 | 742.35 | 2.04 |
|  | Non-social | 4 | 742.58 | 2.27 |
|  | Pro-social+sex | 5 | 743.24 | 2.93 |
|  | Aggression+sex | 5 | 743.28 | 2.97 |
| Testosterone | **Aggression+weight** | **5** | **40.90** | **0.00** |
| (log pg/mL) | Non-social+weight | 5 | 41.19 | 0.29 |
|  | Pro-social+weight | 5 | 41.24 | 0.34 |
|  | Sociosexual+weight | 5 | 41.40 | 0.50 |
|  | Non-social+sex+weight | 6 | 41.58 | 0.68 |
|  | Aggression+sex+weight | 6 | 41.61 | 0.71 |
|  | Pro-social+sex+weight | 6 | 41.66 | 0.76 |
| DHEA | **Aggression** | **4** | **788.56** | **0.00** |
| (pg/mL) | Non-social+sex+weight | 6 | 789.87 | 1.31 |
|  | Sociosexual | 4 | 789.91 | 1.35 |
|  | Non-social+weight | 5 | 790.02 | 1.46 |
|  | Aggression+sex | 5 | 790.07 | 1.51 |
|  | Aggression+weight | 5 | 790.09 | 1.53 |
|  | Sociosexual+weight | 5 | 790.37 | 1.81 |
| Progesterone | **Non-social** | **4** | **686.60** | **0.00** |
| (pg/mL) | Non-social+sex | 5 | 687.67 | 1.07 |
|  | Aggression+non-social | 5 | 688.80 | 2.20 |
|  | Non-social+sociosexual | 5 | 688.84 | 2.24 |
|  | Non-social+age | 5 | 688.86 | 2.26 |
|  | Non-social+weight | 5 | 688.93 | 2.33 |
|  | Non-social+pro-social | 5 | 688.94 | 2.34 |
|  | Behavior frequency models | | | |
| Estradiol | **Aggression** | **4** | **739.63** | **0.00** |
| (pg/mL) | Aggression+sex | 5 | 741.01 | 1.38 |
|  | Aggression+non-social | 5 | 741.52 | 1.89 |
|  | Aggression+age | 5 | 741.71 | 2.08 |
|  | Aggression+pro-social | 5 | 741.78 | 2.15 |
|  | Aggression+weight | 5 | 741.78 | 2.15 |
|  | Aggression+sociosexual | 4 | 741.90 | 2.27 |
| Testosterone | **Pro-social+weight** | **5** | **41.07** | **0.00** |
| (log pg/mL) | Pro-social+sex+weight | 6 | 41.18 | 0.11 |
|  | Sociosexual+weight | 5 | 41.37 | 0.30 |
|  | Aggression+weight | 5 | 41.40 | 0.33 |
|  | Non-social+weight | 5 | 41.42 | 0.35 |
|  | Non-social+sex+weight | 6 | 41.73 | 0.66 |
|  | Aggression+sex+weight | 6 | 41.79 | 0.72 |
| DHEA | **Aggression+non-social+sex+weight** | **7** | **789.85** | **0.00** |
| (pg/mL) | Aggression+sex+weight | 6 | 790.09 | 0.24 |
|  | Aggression | 4 | 790.22 | 0.37 |
|  | Non-social | 4 | 790.23 | 0.38 |
|  | Pro-social | 4 | 790.29 | 0.44 |
|  | Aggression+sex | 5 | 790.34 | 0.49 |
|  | Pro-social+weight | 5 | 790.38 | 0.53 |
| Progesterone | **Non-social+weight** | **5** | **692.46** | **0.00** |
| (pg/mL) | Non-social+sex | 5 | 692.49 | 0.03 |
|  | Non-social+age | 5 | 692.51 | 0.05 |
|  | Non-social | 4 | 692.52 | 0.06 |
|  | Non-social+pro-social+sex | 6 | 693.27 | 0.81 |
|  | Aggression+pros-ocial+sex | 6 | 693.32 | 0.86 |
|  | Non-social+sex+weight | 6 | 693.42 | 0.96 |
